# Gut microbiota characterization in vegetarian adults from the adventist population living in coastal, highland, and jungle regions of Peru

**DOI:** 10.1101/2025.11.26.690875

**Authors:** Pool Marcos Carbajal, Sebastian A Medina-Ramirez, José Luis Yareta Yareta, Miguel Angel Otiniano Trujillo, Mario Salomón Chambi Quispe, Manuel Ruiz Panaifo, Pilar Diaz Rengifo, Cinthia Esther Sias Garay, Alexandra Obregon-Tito, Raul Tito

## Abstract

**Introduction:** The gut microbiota plays a crucial role in various physiological processes, and its composition can be influenced by factors such as diet, genetics, age, and health status. Among these, diet is one of the most significant determinants of microbial balance. A well-balanced diet promotes the growth of beneficial bacteria, reducing intestinal inflammation and the risk of chronic diseases. Vegetarian diets have been proposed to confer beneficial effects on gut health; however, many questions remain regarding microbial profiles, their variability, and the potential influence of geographic and cultural factors—particularly within Latin American contexts. This study aims to address this gap by presenting the first characterization of the gut microbiota in vegetarian adults from the Adventist population living in three distinct regions of Peru.

**Methods:** Stool samples were used as a proxy to analyze gut microbiota composition through 16S rRNA gene sequencing. Standard descriptive analyses were performed to assess bacterial composition, including diversity and relative abundance across samples.

**Results:** The gut microbial communities of Peruvian vegetarians revealed three distinct enterotypes, with distribution varying by region. Enterotype 1 (ET1), predominant in coastal and highland regions, exhibited the highest bacterial richness and diversity. Enterotype 2 (ET2), observed in highland and jungle areas, was characterized by higher levels of *Prevotella*. Enterotype 3 (ET3), more frequent in the jungle region, showed a greater abundance of *Bacteroides* and *Faecalibacterium*.

**Conclusions:** Despite all participants adhering to a vegetarian diet, notable diversity in gut microbiota profiles was observed within this population. While three distinct enterotypes were identified, consistent with findings in other populations, the specific profiles differed from those previously reported. This study highlights the importance of incorporating variables that enable greater resolution in future research, allowing better control of within-population variability, such as that observed in Peruvian vegetarians, and ultimately enhancing the accuracy of microbiome-related conclusions.

## Introduction

The gastrointestinal system is composed of a complex ecosystem where human cells interact with a vast array of microorganisms. It is estimated that the human body harbors bacterial cells in an approximate ratio of 1:1 with human cells [1]. The gut microbiota has garnered increasing attention due to its association with a wide range of diseases, including cancer, neurodegenerative, cardiometabolic, and endocrine disorders [2,3]. Four bacterial phyla: Actinobacteria, Bacteroidetes, Firmicutes, and Proteobacteria, constitute approximately 80–95% of the gut microbiota (4). Among the over 150 species within these phyla, the genera *Bacteroides* and *Prevotella* (Bacteroidetes); *Clostridium, Enterococcus, Lactobacillus*, and *Faecalibacterium* (Firmicutes); and *Bifidobacterium* (Actinobacteria) stand out for their proposed roles in human health [4,5].

The gut microbiota plays active roles in intestinal metabolism. In addition to synthesizing vitamins such as K and B12, neurotransmitters, and enzymes, it contributes critically to the health of colonocytes through the production of short-chain fatty acids (SCFAs), butyrate, propionate, and acetate, via fermentation. It also protects against pathogens through nutrient competition, antimicrobial compound production, and modulation of immune system homeostasis [6]. However, the biological processes associated with the gut microbiota are highly complex, as they are influenced by numerous intrinsic and extrinsic host factors, including genetics, diet, age, medication use, and overall health status [7].

Lifestyle, and more specifically diet, profoundly affects both the composition and function of the gut microbiota [8]. Previous studies have identified three major enterotypes (ET) characterized by the dominance of specific taxa: a diet rich in complex carbohydrates promotes the *Prevotella* enterotype [9], which is typically associated with less industrialized lifestyles and high-fiber diets. In contrast, industrialized lifestyles characterized by prolonged consumption of fats and proteins tend to favor the *Bacteroides* and *Ruminococcus* enterotypes, with the former being more prevalent in urban populations. It has been suggested that microbial communities resembling the *Prevotella* enterotype may be more beneficial to health than those resembling the *Bacteroides* or *Ruminococcus* enterotypes, which have been associated with chronic conditions such as diabetes and hypertension, commonly referred to as “diseases of civilization” [10,11].

On the other hand, according to the Academy of Nutrition and Dietetics of the United States, a well-planned vegetarian or vegan diet can be appropriate for individuals at all stages of life and may offer health benefits [12], such diets can enhance nutrient absorption and promote the growth of beneficial bacterial genera such as *Clostridium, Lactobacillus, Faecalibacterium*, and *Prevotella*, while potentially reducing the abundance of genera like *Bacteroides* and *Bifidobacterium*, leading to a gut microbial community with anti-inflammatory properties[13,14].

In Peru, as in much of Latin America, high consumption of processed foods is a key factor contributing to elevated obesity rates [15,16]. Nevertheless, a small but growing segment of the population follows a diet rich in fruits and vegetables, with its distribution varying across regions and shaped by cultural and socioeconomic factors [17]. However, the impact of vegetarian and vegan diets on gut microbiota composition and metabolic profiles within Latin American populations remains largely understudied [18].

This knowledge gap is further compounded by the region’s high ecological and cultural diversity [19], making it essential to conduct region-specific characterizations before broader generalizations can be made. In this context, the present study aims to address this gap in Peru by characterizing the gut microbiota composition in vegetarian adults from the Adventist population living in cities across three distinct regions of the country. The findings are intended to contribute to a better understanding of how vegetarian diets may influence microbiological health within culturally and ecologically diverse settings.

## Methods and Study Population

A descriptive cross-sectional study was conducted between March and December 2022 among vegetarian individuals aged 18 to 70 years, residing in cities located in the three main geographic regions of Peru. The target population consisted of members of the Seventh-day Adventist community in Peru, distributed across various localities. The selected cities included: Trujillo, Tacna, and Chiclayo (Coastal region); Puno, Cusco, and Arequipa (Highland region); and Pucallpa, Loreto, San Martín, and Puerto Maldonado (Jungle region).

Participants were recruited using non-probability snowball sampling within the Adventist community, given the geographic dispersion of the population and the absence of a centralized registry that would allow for probabilistic sampling. Individuals were excluded if they had chronic diseases, a history of inflammatory gastrointestinal conditions, recent use of corticosteroids or antibiotics, or had taken dietary supplements (probiotics, prebiotics, or synbiotics) in the previous six months. Detailed inclusion and exclusion criteria are provided in **Supplementary Material 1**.

A vegetarian was defined as any individual who reported not consuming red or white meat or fish on a regular basis for at least the past 12 months, and who also self-identified as vegetarian. Participants were considered apparently healthy if they did not meet any of the exclusion criteria.

### Procedures

Interested volunteers were individually contacted and informed about the study. After an initial screening process, eligible participants were invited to attend a medical check-up. If inclusion criteria were met, a biological sample was collected. Samples were obtained at the healthcare center closest to the participant’s place of residence by trained healthcare personnel.

### Clinical Data and Biological Sample Collection

#### Study Variables

Anthropometric measurements were taken from all participants using a fixed stadiometer (SECA® model 213; precision: 0.1 cm) to measure height, and a digital scale (INDRA®; precision: 0.1 kg) to measure weight. Body mass index (BMI) was calculated as weight in kilograms divided by height in meters squared and was used to classify nutritional status according to the World Health Organization (WHO) criteria [19]. For older adults, adjusted cut-off points were applied, as provided by the official form available at: https://alimentacionsaludable.ins.gob.pe/adultos-mayores-0. Additionally, a fasting blood sample (minimum 8 hours) was collected from each participant to determine levels of glucose, triglycerides, cholesterol, hemoglobin, and hematocrit. Stool samples were collected in sterile containers and immediately frozen at –20°C for later processing. Each stool sample was divided into two aliquots: one preserved for nucleic acid extraction and sequencing, and the other used for parasitological analysis.

Anthropometric measurements and biological sample collection were carried out by previously trained members of the research team with a background in healthcare.

### DNA Extraction and 16S rRNA Gene Sequencing of Microbial RNA

Total DNA extraction was performed in duplicate using the QIAamp DNA Stool Mini Kit, following the manufacturer’s instructions, at the Molecular Biology Research Laboratory (LIBM) of UpeU. Extracted DNA was quantified using the Qubit 1X dsDNA BR Assay Kit. For subsequent molecular analyses, only the sample with the highest DNA concentration was used. The 16S rRNA gene of microbial RNA was sequenced using amplicon sequencing with paired-end reads (read length: 301 bases), employing the Herculase II Fusion DNA Polymerase and the Nextera XT Index V2 Kit on the Illumina MiSeq platform (Macrogen). The resulting sequences were processed using the LotuS [20] and DADA2 [21] pipelines to generate amplicon sequence variants (ASVs). Taxonomic assignment of ASVs was performed using the SILVA database (version 138, SLV_nr99_v138.1). Taxonomic levels from domain to genus were assigned using the assignTaxonomy function from the DADA2 package in R. To prevent issues related to unassigned taxonomic labels, any ASVs lacking classification at a given taxonomic level were labeled with the prefix “uc” (unclassified), followed by the last assigned level.

Prior to downstream analyses, sequences annotated as belonging to chloroplast class, mitochondrial family, archaea, and unknown bacteria were removed. The phyloseq (v1.36.0) and MicroViz (v0.11.0) packages in R were used for data curation and figure generation.

### Statistical Analysis

To evaluate differences between groups, sequencing data were aggregated at the genus taxonomic level, and the read count was rarefied to 7,600 reads per sample. Differential abundance analysis was performed using non-parametric tests: the Kruskal–Wallis rank-sum test for comparisons involving more than two groups, and the Wilcoxon rank-sum test for comparisons between two groups. P-values were adjusted using the Benjamini–Hochberg correction, and values below 0.05 (adjusted p-value, adjP) were considered statistically significant. Enterotyping (community typing) was performed using a Dirichlet multinomial mixture model in R, as previously described [22], based on the genus-level abundance matrix. The optimal number of Dirichlet components was determined to be three, according to the Bayesian Information Criterion [22]. The distribution of enterotypes across regions was analyzed using Pearson’s chi-squared test (*p < 0*.*05*).

### Ethical Considerations

The study was approved by the Ethics Committee of the Universidad Peruana Unión (UPeU), under resolution No. 2022-CEUPeU-1294. Written informed consent was obtained from each participant prior to enrollment.

## Results

A total of 86 participants were recruited, of whom two were excluded, one due to missing clinical history data and another due to potential sample contamination, resulting in a final sample of 84 individuals. Of the participants, 41.7% were male, with a mean age of 38 years. Regarding clinical results, hemoglobin levels were highest in the highland region compared to the jungle and coastal regions (*15*.*9 mg/dL, 13*.*2 mg/dL, and 14*.*1 mg/dL, respectively*). BMI was similar across the three regions and fell within the normal range. No significant differences were observed among regions in sex distribution, age, glucose, cholesterol, or triglyceride levels (**Table 1**).

**Table 1.** Clinical characteristics of the study population.

| Variable | Total | Coastal | Highland | Jungle | adjP |
| --- | --- | --- | --- | --- | --- |
| N | 84 | 22 | 27 | 35 |  |
| Male sex, n(%) | 35 (41.7) | 11 (50.0) | 10 (37.0) | 14 (40.0) | 0.848 |
| BMI (median, [IQR]) | 22.53 [4.12] (16.2-32.32) | 23.38 [4.75] (16.2-32.32) | 22.6 [4.15] (17.63-29.03) | 21.78 [3.25] (16.36-28.58) | 0.547 |
| Age (median, [IQR]) | 36.5 [21] (12-86) | 37 [23.25] (15-86) | 39 [21.5] (14-67) | 35 [15] (12-70) | 0.946 |
| Hemoglobin, g/dL (median [IQR]) | 13.3 [2.97] (7.8-18.3) | 13.5 [2.53] (7.8-17.9) | 13.7 [2.5] (10.9-18.3) | 12.87 [1.66] (10.1-16.66) | 0.004 |
| Glucose, mg/dL (median [IQR]) | 80 [14.45] (49.9-119.2) | 82.7 [14.28] (59.9-119.2) | 82 [18.2] (49.9-106.8) | 79 [10.85] (55.1-96) | 0.250 |
| Cholesterol, mg/dL (median [IQR]) | 160.5 [53.3] (84-262) | 158.9 [41.18] (100.1-239.6) | 172.3 [51] (102.9-240) | 150.6 [41.5] (84-262) | 0.946 |
| Triglycerides, mg/dL (median [IQR]) | 109 [60.7] (40.2-400) | 99.7 [94.88] (40.2-302.1) | 104 [56.35] (43.3-400) | 117 [48.4] (50-242) | 0.197 |

adjP = p-value adjusted using the Benjamini-Hochberg correction. IQR = interquartile range. BMI = body mass index. Values in bold indicate statistical significance (adjP < 0.05).

The structure of gut microbiota communities varied significantly across the Peruvian regions (coast, highlands, and jungle) (Adonis test R^2^ = 0.06, p = 8E-04), as shown in **Figure 1a**, although Principal Coordinates Analysis (PCoA) did not reveal a distinct clustering of samples by region. Thirteen bacterial genera were identified with differential abundance between regions (**Figure 1b**, Kruskal– Wallis test, adjP < 0.05). In samples from coastal cities, the dominant bacterial genera included *Clostridium sensu stricto 1, Intestinibacter, Romboutsia*, and *UCG*.*003*. In contrast, samples from the jungle region showed higher relative abundances of *Sutterella, UCG*.*004, UCG*.*010*, and *Lachnospira*, while the highland region displayed low abundance of *UCG*.*003*.

**Figure 1.**
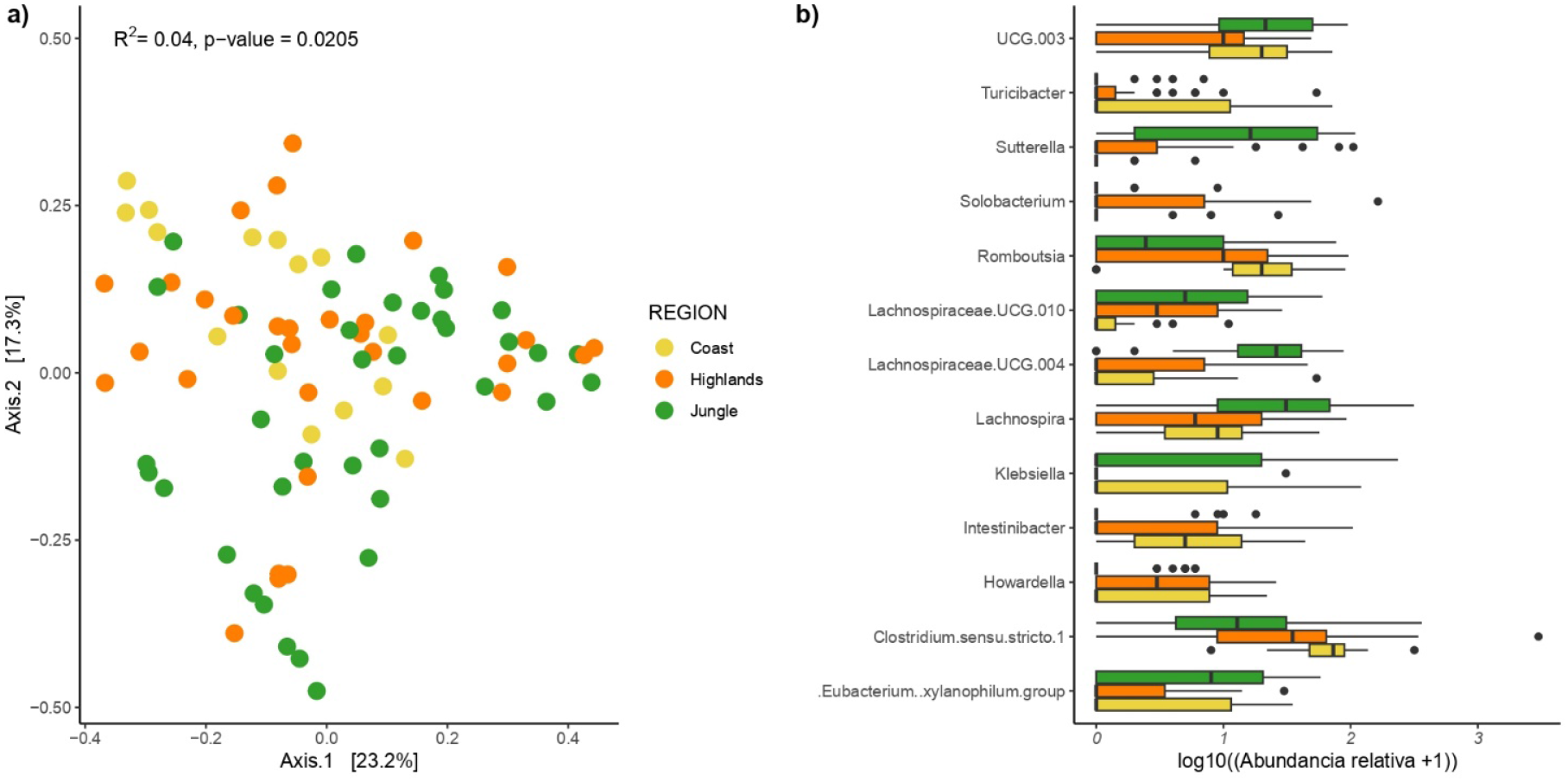
Composition of gut microbial communities in Peruvian vegetarians by geographic region. Principal Coordinates Analysis (PCoA) of community structure by region (a) and bacterial genera showing differential relative abundance between regions (b). Kruskal–Wallis test, adjP < 0.05.

After clustering microbial communities based on compositional similarity, the 84 microbiota profiles were grouped into three constellations or enterotypes: ET1, ET2, and ET3 (**Figure 2a**). The distribution of enterotypes differed significantly by geographic region (Pearson’s chi-squared test, p < 0.05), with the coastal region showing no presence of ET3 and a higher proportion of ET1 (**Figure 2b)**.

**Figure 2.**
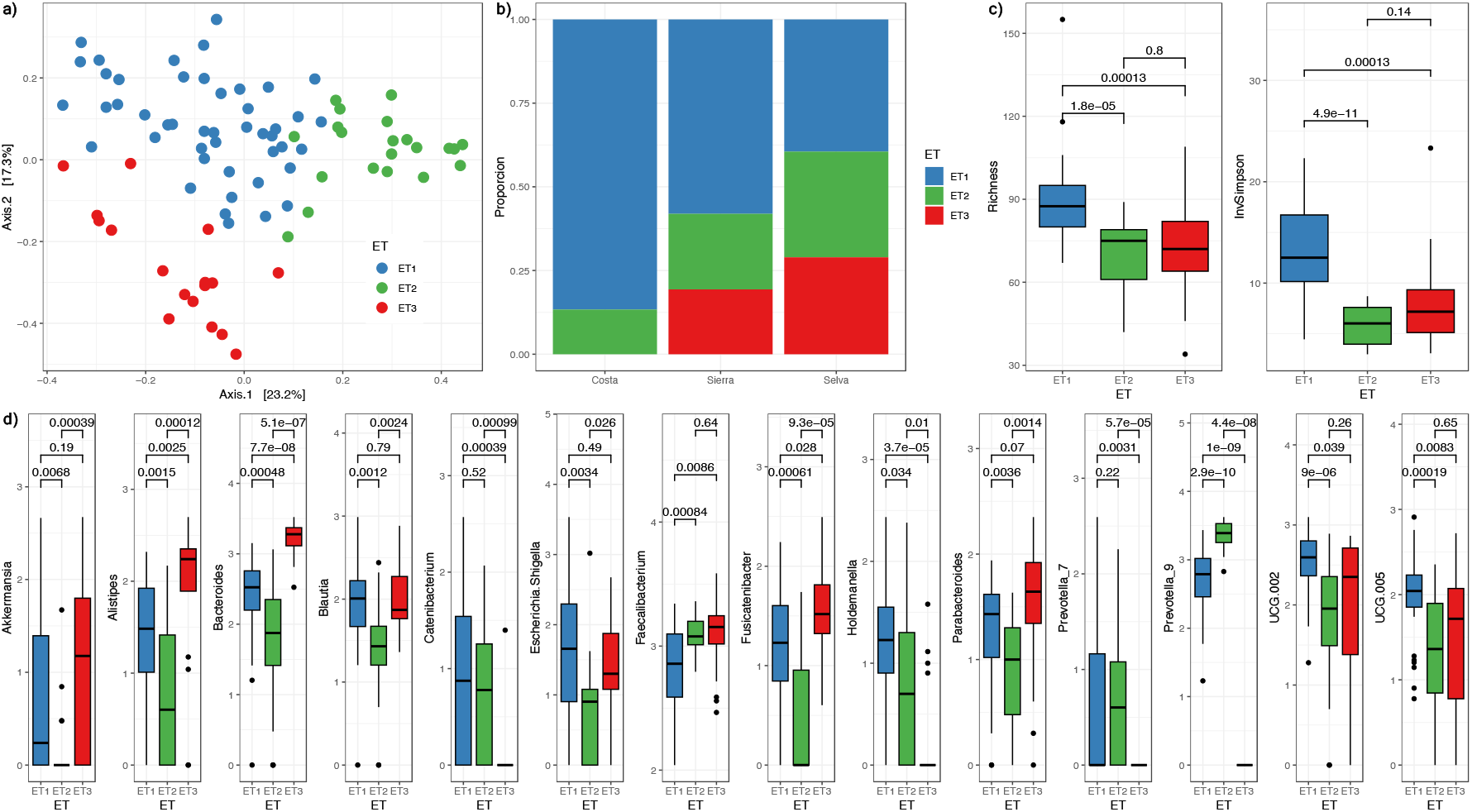
Enterotypes in the Peruvian vegetarian population. (a) Principal Coordinates Analysis (PCoA) showing three clearly identified enterotypes. (b) Geographic distribution of enterotypes, highlighting the absence of ET3 in coastal participants. (c) Comparison of microbial richness and diversity across enterotypes (ET1, ET2, ET3), including richness (“Richness”) and inverse Simpson index (“InvSimpson”), with ET1 displaying significantly higher diversity than the other two enterotypes. Bars indicate significant differences with corresponding p-values. (d) Relative abundance of the most frequent bacterial genera across the three Peruvian enterotypes identified in this cohort. Thirty genera showed statistically significant differences in abundance among enterotypes (Kruskal–Wallis test, adjP < 0.05, followed by Wilcoxon post-hoc test).

ET1 exhibited significantly greater microbial richness and diversity compared to ET2 and ET3 (p < 0.05), as measured by both species richness and the inverse Simpson index, which incorporates abundance-weighted diversity. No significant differences were observed between ET2 and ET3 in these diversity metrics (p > 0.05) (**Figure 2c**). A total of 30 bacterial genera showed differential relative abundance across the three Peruvian enterotypes (Kruskal–Wallis test, adjP < 0.05), considering only genera with a prevalence above 20% in the total sample.

ET1 was characterized by the lowest abundance of *Faecalibacterium* and the highest abundances of *UCG-002* and *UCG-005*, compared to ET2 and ET3. ET2 showed a higher abundance of *Prevotella*, alongside the lowest abundances of Escherichia/Shigella, Akkermansia, Blautia, and Parabacteroides. In contrast, ET3 displayed higher levels of *Bacteroides, Fusicatenibacter*, and *Alistipes*, and lower abundances of *Holdemanella* and *Catenibacterium*.

## Discussion

Our study reveals differences in the gut microbiota of vegetarians from the three natural regions of Peru (coast, highlands, and jungle). As expected, hemoglobin levels were higher among participants from the highlands compared to those from the coast and jungle, while no differences were observed in other clinical markers assessed across these populations. [23].

Analysis of gut microbiota composition revealed considerable variability within the Peruvian vegetarian community, identifying three distinct enterotypes with regionally differentiated distributions. While previous studies have reported enterotypes associated with lifestyle and, consequently, dietary patterns, our findings show the presence of distinct enterotypes within a population sharing a similar diet (i.e., vegetarian), and notably, these enterotypes were not dominated by *Prevotella*, the genus typically expected to predominate in such populations based on prior research [17,18]. These findings suggest that the broader macroenvironment in which individuals reside may influence gut microbiota composition to a degree that is not overridden by diet alone. Therefore, it is critical for human microbiome studies to incorporate comprehensive metadata and to emphasize the inclusion of diverse populations, even when the study population, such as Peruvian vegetarians, might initially be assumed to be homogeneous. [17,24].

Hemoglobin levels observed in the highland population (13.7 g/dL) reflect a well-documented physiological adaptation to reduced ambient oxygen levels with increasing altitude and decreasing atmospheric pressure. This elevation in hemoglobin facilitates improved oxygen transport, helping to counteract environmental hypoxia. This association has been primarily documented in non-vegetarian populations, while vegetarian populations are often considered to be at increased risk for anemia [25,26]. However, in our study, no participants from any of the three regions had mean hemoglobin levels consistent with anemia.

Other clinical markers, including BMI, glucose, cholesterol, and triglycerides, showed similar distributions across the three regions. These findings suggest that no major regional differences exist in these clinical parameters.

In contrast to the clinical profile, which was largely uniform across regions (with the exception of hemoglobin), the gut microbiota composition differed markedly. These differences may reflect the host’s habitat and ecological niche. For example, coastal participants showed higher relative abundances of Clostridium sensu stricto 1, Intestinibacter, and Romboutsia, while participants from the jungle exhibited greater abundances of Sutterella, Lachnospira, and UCG-004. Although exploring specific associations between these taxa and individual dietary elements lies beyond the scope of this study, we recognize that regional differences in the availability of fruits and vegetables, exposure to pesticides, endemic microorganisms, and water quality may significantly influence gut microbial composition [27]. These findings underscore geography as an influential variable in shaping gut microbiota, even within a relatively homogeneous dietary group such as vegetarians.

Gut enterotypes, defined as distinct and stable clusters of microbial community profiles, have been reported across multiple human populations [28]. In contrast to previous studies suggesting that fiber-rich, low-meat diets are associated with enterotypes dominated by *Prevotella* [29], our study identified three distinct enterotypes with region-specific distributions, only one of which was characterized by a high abundance of *Prevotella* [30] These findings differ from the model proposed by Arumugam et al., who conducted a large phylogenetic mapping study and described three main enterotypes, dominated respectively by *Bacteroides, Prevotella*, and *Ruminococcus* [29].

Enterotype 1 (ET1), characterized by higher microbial richness and diversity as well as the presence of UCG-002 and UCG-005, was detected across all three regions but was far more prevalent in the coast and highlands, where it represented the majority of participants. This enterotype may be associated with more stable gut ecosystems and potentially with greater resilience to gastrointestinal disorders. In contrast, Enterotype 2 (ET2), which showed a higher abundance of Prevotella—a genus typically linked to fiber-rich diets commonly found in traditional agrarian societies—was dominant in the jungle region and present at lower proportions in the highlands and even less so on the coast. Since Prevotella-dominated enterotypes have been previously associated with traditional diets, its limited presence on the coast and higher frequency in more remote regions suggest that Prevotella dominance may be linked to a specific subtype of plant-based diet, rather than broadly representing all high-fiber, low-animal-product diets. [31,32]

Enterotype 3 (ET3), which was absent in coastal participants but predominant in the jungle and partially represented in the highlands, was characterized by higher relative abundances of *Bacteroides, Fusicatenibacter*, and *Alistipes*, taxa commonly associated with omnivorous dietary patterns. This apparent mismatch between microbial profile and the self-reported vegetarian diet of participants from the highlands and jungle suggests that previous associations between these taxa and omnivorous diets may not be sufficiently specific. It is possible that these profiles are shaped by particular characteristics of the vegetarian diet or even by food preparation methods rather than by food categories alone. There is no doubt that the taxonomic diversity and differential abundance of certain genera across enterotypes have important functional implications. The higher levels of *Faecalibacterium* in ET1 and lower levels of *Prevotella* in ET2 suggest differing metabolic capacities among these communities, which in turn may impact host health.

In Peru, evidence regarding gut microbiota in this population is scarce. However, previous studies in pediatric populations with anemia have reported high abundance of Prevotella, which partially aligns with our findings in participants from the jungle. This suggests that diet alone may not be the sole factor influencing microbial composition [33,34].

### Limitations

The main limitation of this study is the sample size. While it appears sufficient for an initial characterization and allowed the detection of significant variability, it may still be insufficient to fully capture the diversity of gut microbial communities across Peru’s different regions. Another limitation is the descriptive nature of the study, which restricts the ability to establish causal relationships or to explore the complexity of microbial interactions in depth.

Finally, given the heterogeneity of human populations in countries with such megadiverse natural and cultural environments, it is critical to ensure that study samples are representative of the broader population to which the findings are expected to be generalized.

### Strengths

The main contribution of this study lies in its examination of variability within a seemingly homogeneous population—Peruvian vegetarians—which, upon molecular analysis, reveals substantial heterogeneity that might be overlooked if assessments focused solely on clinical markers. Our study also contributes to expanding knowledge about the human gut microbiome, which is currently centered largely on European, Asian, or North American populations [35].

Despite ongoing debate around the use of enterotypes, the analysis of these community clusters offers a useful tool to identify objective differences between regions, thereby contributing to a deeper understanding of microbiota variability within populations. Additionally, this study provides evidence for the potential impact of non-dietary factors on gut microbiota composition, offering a foundation for future confirmatory research on the social and ecological determinants of the human microbiome.

In conclusion, this study highlights previously undescribed variability in the gut microbiota of vegetarian communities. The three identified enterotypes—clusters defined by specific taxonomic patterns—were distributed differently across geographic regions, revealing profiles that align with previously described enterotypes associated with high-fiber, plant-based diets as well as those commonly linked to omnivorous diets, despite none of the participants reporting animal product consumption. These observations emphasize two critical points: a) The gut microbiota of seemingly homogeneous populations, such as Peruvian vegetarians, may harbor hidden variability, which could lead to misleading conclusions if not properly addressed; b) Regional factors—including geography, water quality, exposure to various foodborne pesticides, and methods of food preparation—may significantly contribute to the observed microbiota variation. Therefore, rigorous control of confounding variables, based on comprehensive metadata, is essential for establishing causal relationships that may lead to clinically relevant applications of microbiome research.

## Author Contributions

All authors declare that they meet the authorship criteria recommended by ICMJE

## Conflict of Interest

The authors declare no conflict of interest related to the publication of this article.

## Funding

This study was funded by Universidad Peruana Unión

## Notes

**Conflicts of Interest** The authors declare no conflicts of interest regarding this manuscript.

### Competing Interest Statement

The authors have declared that no competing interests exist.

